# Elucidating directed neural dynamics of scene construction across memory and imagination

**DOI:** 10.1101/2025.07.22.666182

**Authors:** C. Kindler, J. Taube, P. Leelaarporn, R. Stirnberg, C. McCormick

**Author notes:** Corresponding author: Cornelia McCormick Department for Old Age Psychiatry and Cognitive Disorders Universitätsklinikum Bonn Venusberg-Campus 1 Gebäude C.82 D-53127 Bonn.

## Abstract

Autobiographical memory (AM) and imagination both rely on the brain’s ability to construct vivid, coherent mental representations of past or imagined experiences. A central cognitive process underlying these functions is scene construction—the mental generation of spatially organized, imagery-rich representations of environments. Using ultra-high field 7T fMRI combined with Dynamic Causal Modeling (DCM), we examined directed effective connectivity among key nodes of the default mode network: the ventromedial prefrontal cortex (vmPFC), hippocampus, and precuneus during object imagery, single-scene construction, and AM retrieval. Our results revealed distinct patterns of network dynamics depending on task demands. During AM retrieval, effective connectivity was characterized by vmPFC-driven top-down modulation primarily targeting the precuneus, supporting the temporal and self-relevant structuring of episodic memory. In contrast, extended scenario imagination engaged hippocampal bidirectional influences with both vmPFC and precuneus, reflecting the dynamic simulation and integration of unfolding events. Single Scene construction shifted network leadership to the precuneus, which exerted modulatory effects on both vmPFC and hippocampus, consistent with its pivotal role in spatial integration and the generation of coherent mental scenes. Object imagery showed minimal stable connectivity within this network, suggesting limited engagement of the hippocampal-default mode circuit during simpler visual representations. Together, our findings highlight a flexible, task-dependent reorganization of effective connectivity within the vmPFC– hippocampus–precuneus network during the construction of rich mental experiences. Scene construction emerges as a unifying mechanism linking memory and imagination, with the direction and strength of neural interactions adapting according to whether temporal or spatial demands predominate.

**Highlights:**

- Autobiographical memory is reflected by vmPFC-driven effective connectivity
- Effective connectivity varies flexibly amongst vmPFC, hippocampus and precuneus
- Scene construction emerges as a core process across memory and imagination

## Introduction

Autobiographical memory (AM) and imagination are both grounded in the capacity to construct imagery-rich mental representations of past or fictitious experiences. A central cognitive process underlying this capacity is scene construction, the ability to mentally generate coherent, spatially organized representations of a setting or situation (Hassabis et al., 2007; Maguire & Mullally, 2013). This mechanism is thought to serve as a phenomenological glue linking memory and imagination, due to their shared reliance on vivid, immersive, and flexible mental experiences (Hassabis et al., 2007; Maguire & Mullally, 2013).

Neuroimaging studies have consistently shown that both AM and imagination engage a distributed neural network, especially focusing on the hippocampus, ventromedial prefrontal cortex (vmPFC), and precuneus (see also default mode network (DMN), Svoboda et al., 2006; Hassabis & Maguire, 2009; Bellana et al., 2017). These regions are not only co-activated during task performance but are thought to support integrative functions central to internally generated cognition, such as perspective-taking, self-relevance, and temporal structuring (Spreng & Grady 2010; McCormick et al., 2015).

While functional connectivity studies support overlapping recruitment of this network in AM and imagination, they do not reveal the directionality of interactions, that is, how information flows between regions. Effective connectivity approaches, such as Dynamic Causal Modeling (DCM, Friston et al. 2011; Zeidman et al. 2019), provide a more precise framework for addressing this question. Prior work using DCM and related methods has shown that during AM, vmPFC often drives hippocampal and posterior cortical activity, suggesting a top-down control mechanism (Jacques et al., 2011; Nawa & Ando, 2019; McCormick et al., 2020). However, whether this pattern generalizes to novel or future-oriented imagination remains unclear.

Scene construction is central to both AM and imagination, yet the underlying network dynamics may differ. Constructing a single static scene may rely on different neural mechanisms than integrating multiple scenes into temporally extended scenarios, as required for imagining future events (McCormick et al. 2018a; Taube et al., 2025 preprint). The hippocampus has been implicated in both static and dynamic forms of construction, while the vmPFC appears increasingly engaged as scenarios become more elaborate and temporally extended (Barry et al., 2019; Monk et al., 2021). In addition, the precuneus has been implicated in self-centered visual imagery (Cavanna & Trimble, 2006; Dadario et al., 2023).

In the present study, we address these questions by examining effective connectivity patterns during AM retrieval and three forms of imagination: object construction, single-scene construction, and extended scenario construction. Using 7T fMRI and DCM, we examined directed interactions between vmPFC, hippocampus, and precuneus across two independent datasets. We hypothesized that AM would replicate previously observed vmPFC-driven influences, and that extended scenario construction would share some important network dynamics, albeit potentially adding multidirectional influences. Further, we hypothesize that scene construction underlies more hippocampal-driven dynamics. We included object construction as a control condition, expecting that effective connectivity measures would not show stable results.

By clarifying the dynamic interplay between key nodes in the DMN, this study seeks to refine our understanding of the neural architecture of constructive cognition, and to shed light on how the brain flexibly generates coherent, mental representations from memory and imagination.

## Materials and methods

### Data sources

In the present study, we analyzed data from two previously conducted experiments: one published dataset (Leelaarporn et al., 2024, accessible upon request.) and one dataset currently published as a preprint (Taube et al., 2025 preprint). Both studies employed the same 7 Tesla fMRI acquisition protocol and involved young, healthy adult participants. While the experimental design was similar across both studies, the first dataset focused on AM retrieval, while the second examined visual imagination. Given the methodological consistency and complementary cognitive domains, these datasets provide a suitable foundation for the current comparison of effective connectivity profiles.

A total of 43 participants were included: 23 from the AM study (mean age = 26.4 ± 4.1 years; 12 female, 11 male) and 19 from the imagination study (mean age = 27.9 ± 3.7 years; 11 female, 8 male). All participants were right-handed, had normal or corrected-to-normal vision, no history of neurological or psychiatric disorders, and had completed at least 12 years of formal education. Informed written consent was obtained from all participants in accordance with the respective institutional ethics approvals.

In both studies, participants engaged in covert, imagery-based tasks in response to verbal cues, followed by vividness ratings. The AM task involved the retrieval of personally experienced events during fMRI scanning: Participants completed 20 randomized trials of AM retrieval and 20 trials of a simple mental arithmetic (MA) baseline task, which was adapted from McCormick et al. (2015) and also employed in Leelaarporn et al. (2024). Each trial lasted up to 17 seconds, with a jittered interstimulus interval of 1–4 seconds. During AM trials, participants were presented with event cues (e.g., “birthday celebration”) and instructed to select a personal event that was specific in time and place and had occurred more than one year ago. Once selected, they pressed a button and were then instructed to vividly recall the event in as much detail as possible. In MA trials, participants solved basic arithmetic problems (e.g., 13 + 53) and then iteratively added 3 to the solution. After each AM trial, vividness was rated; after each MA trial, task difficulty was rated.

The imagination conditions included mental construction of isolated objects, static scenes, or dynamic scenarios unfolding over time (Taube et al. preprint). In short, the experiment was separated into 1. an extensive training session, during which participants learned about the different imaginative and baseline conditions, 2. an eye-tracking experiment and 3. the main fMRI experiment. During the experiment, participants performed three types of mental imagery and a baseline condition. In the *object imagery* (OB) condition, participants visualised a single, detailed item in isolation, akin to a product photograph (e.g., an espresso). The scene imagery (SC) condition required mentally constructing a static, spatially coherent environment containing multiple elements, like a static postcard view, without imagined motion (e.g., a street café). In the scenario/event (SCN) imagery condition participants generated dynamic mental simulations that unfolded over time, incorporating a sequence of events and settings (e.g., meeting a friend for coffee). Participants were instructed to create novel mental images without relying on prior memories. The baseline condition required participants counting the number of characters in meaningless letter strings, providing a cognitively demanding control condition without imagery. While all imagery conditions engaged visual mental simulation, they were designed to vary systematically in spatial and temporal complexity. During fMRI scanning, 45 imaginative (15 per condition) and 15 baseline trials were randomly interleaved and presented for 10 sec, followed by a self-paced vividness rating for 5 sec.

### Functional MRI data acquisition and preprocessing

For the purpose of the current study, we re-analysed the fMRI data in order to examine effective connectivity between key brain regions. Importantly, in both datasets, the identical fMRI sequence on the same MAGNETOM Plus 7 Tesla MRI Scanner (Siemens) was used: a custom interleaved multishot 3D echo planar imaging (EPI) sequence (Stirnberg & Stöcker, 2021) with the following parameters: TE = 21.6 ms, TR_volume_ = 3.4 s, flip angle = 15°, 6/8 partial Fourier along the primary phase-encode direction, and an oblique-axial slice orientation aligned with the anterior-posterior commissure line. The matrix size was 220 × 220 × 140, providing whole-brain coverage at 0.9 mm isotropic resolution, which mitigates intra-voxel dephasing in brain areas particularly prone to inhomogeneities of the MRI main magnetic field such as the ventromedial prefrontal cortex.

To achieve high spatiotemporal resolution across the whole brain with sufficient signal-to-noise ratio (SNR) and BOLD-optimal TE at 7T, several advanced sequence features were combined:

A. Skipped-CAIPI 3·1×7_z2_ sampling, which uses SNR-optimized 7-fold CAIPIRINHA undersampling and interleaved 3-shot segmentation with online 2D GRAPPA reconstruction (Stirnberg & Stöcker, 2021). (B) One externally acquired phase correction scan per volume rather than the typical per-shot correction (Stirnberg & Stöcker, 2021). (C) Variable echo train lengths, skipping only the latest EPI echoes outside a semi-elliptical k-space mask defining 0.9 mm voxel resolution (Stirnberg et al., 2017). (D) Rapid slab-selective binomial-121 water excitation replacing time-consuming fat saturation (Stirnberg et al., 2016).

For both datasets two main functional sessions (∼15 minutes each) were acquired. Each session included up to 264 volumes, with the first five images (covering a 17-second waiting period before task onset) excluded to avoid non-steady-state signals. The session concluded with a standard 3 mm isotropic two-echo gradient-echo field-mapping scan (35 seconds) and acquisition of a T1w image (0.6 mm isotropic whole-brain multi-echo MP-RAGE, see Leelaarporn et al. 2024 or Taube et al. 2025, preprint for more specifics).

Both datasets were also preprocessed identically using the SPM12 (Statistical Parametric Mapping) software package (www.fil.ion.ucl.ac.uk/spm/) running on MATLAB R2017a (MathWorks; https://matlab.mathworks.com/). Anatomical and functional images for each participant were reoriented to align with the anterior–posterior commissure axis. Field maps (phase and magnitude images) were used to compute voxel displacement maps (VDMs), which were applied during realignment and unwarping to correct for geometric distortions in the EPI images. The averaged anatomical scans were co-registered to the functional images. Subsequently, motion-corrected and co-registered functional data were normalized to Montreal Neurological Institute (MNI) space and spatially smoothed with a 6 mm full-width at half-maximum (FWHM) Gaussian kernel.

### Subject-Level DCM Analysis

Dynamic Causal Modeling (DCM) is a Bayesian framework for inferring directed (effective) connectivity between brain regions, allowing researchers to model how activity in one region influences activity in another under different experimental conditions (Friston et al. 2011). The current DCM analysis was conducted separately for the AM dataset and, for the scene-based imagination dataset, separately for the SCN, SC, and OB conditions.

#### Regions-of-interest selection and time series extraction

To keep our models simple and focused on the main hypotheses, we specified three regions of interest (ROIs) in the left hemisphere: the ventromedial prefrontal cortex (vmPFC; [-2, 57, -14]), hippocampus ([-27, -12, -26]), and precuneus ([-2, -64, 33]), using 5 mm spherical masks centered on peak voxels identified in the AM > MA contrast from Leelaarporn et al. (2024). These coordinates also showed task differences in the imagery task, and they are consistent with regions commonly reported in the AM literature (Svoboda et al. 2006). While AM-related fMRI activation is typically bilateral, prior research suggests a slight left-hemisphere dominance (Maguire et al. 2001). Although an ideal model would include a more extensive set of bilateral ROIs, we limited our selection to the left hemisphere to maintain tractability and interpretability within the DCM framework. For each ROI, the summary time series (the first eigenvariate) of all voxels within each ROI was extracted using F-contrasts: AM condition versus the baseline MA condition (AM dataset) and SCN, SC and OB versus the baseline NW condition (scene-based imagination dataset). The first eigenvariate was extracted, as commonly done in DCM analysis, to maximize the signal-to-noise ratio and reduce dimensionality.

#### Model Specification

Our main goal was to illuminate similarities and differences during AM and scene-based imagination in effective connectivity between key brain nodes involved in these tasks. Thus, we specified four models with systematically varying directional connections (i.e., modulatory effects in the B-matrix). The first model was a nearly fully connected model but restricted to the connections hypothesized to be as modulated by the AM and /or imagery conditions while self-connections were left unmodulated (Fig 1A). In the second model, the condition modulated top-down connections: vmPFC → hippocampus, vmPFC → precuneus, and hippocampus → precuneus (Fig 1B). The third model was restricted to hippocampus-driven modulations: hippocampus → vmPFC and hippocampus → precuneus connections (Fig. 1C). In the fourth model, the condition modulated the precuneus → hippocampus, precuneus → vmPFC, and hippocampus → vmPFC connections (Fig. 1D). In all models, self-connections in the B-matrix were not specified as they were not relevant to the specific research question. In addition, in all models, each task conditions (AM, SCN, SC, or OB) and the respective baseline condition (MA or NW) influenced all regions.

**Figure 1.**
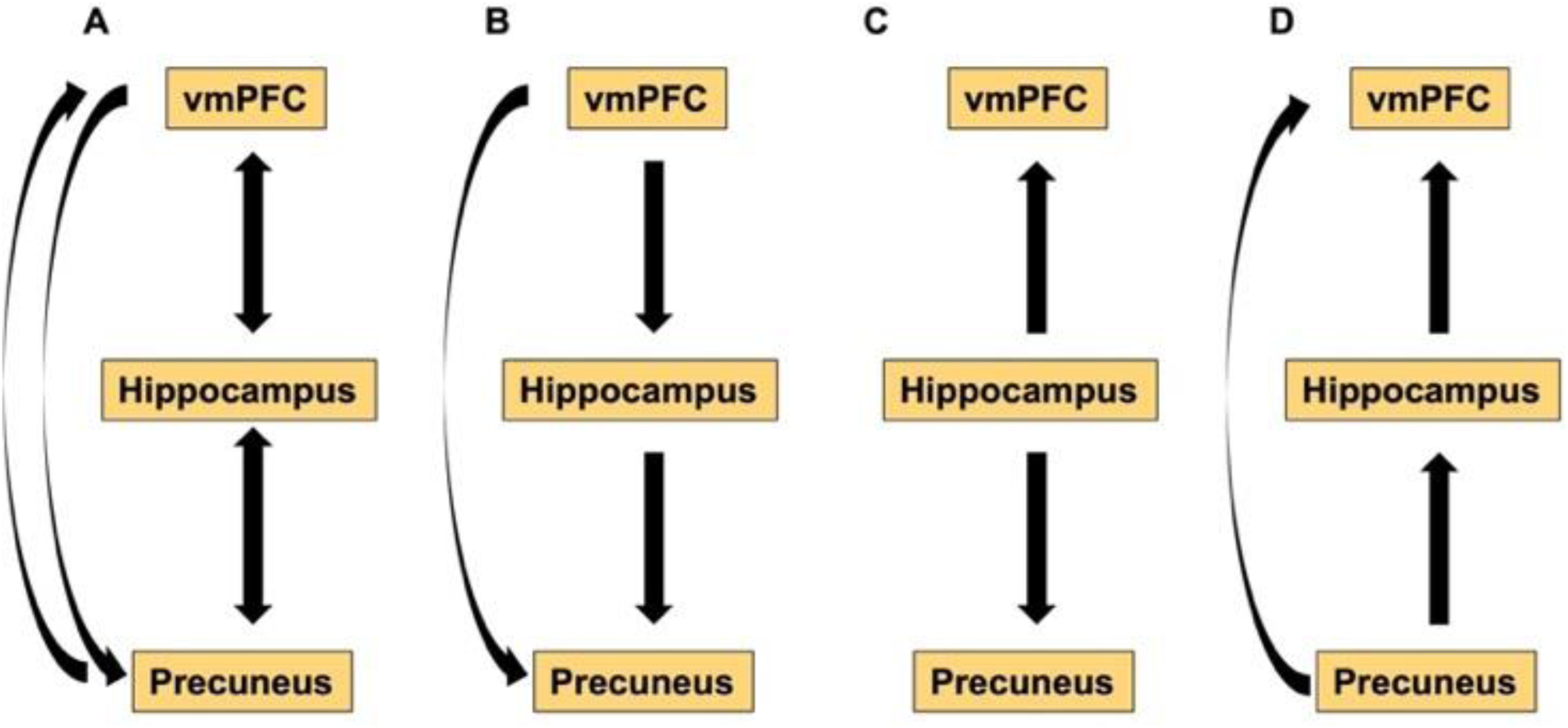
Model Specifications for the vmPFC–Hippocampus–Precuneus Networks. Schematic overview (A–D) illustrating the hypothesized directional influences among vmPFC, hippocampus, and precuneus as set up in each model specification. Arrows represent proposed modulated connections and their direction of effect.

### Group-Level DCM Analysis

Modulatory effects of the experimental conditions were assessed using a hierarchical Parametric Empirical Bayes (PEB) framework, modelling only the B-matrix parameters (Friston et al., 2016). To identify the most plausible model architecture across participants, we first conducted random-effects Bayesian model selection (RFX-BMS) among four competing models within each cognitive condition. RFX-BMS accounts for inter-individual variability and estimates how frequently each model occurs in the population (Stephan et al., 2009). Model comparison was based on several complementary metrics. The expected posterior probability (EPP) indicates how often a model would be selected across the population, while the exceedance probability (XP) quantifies the likelihood that a given model is more frequent than any other in the set. The protected exceedance probability (PXP) further adjusts for the possibility that observed differences in model frequency arose by chance. To evaluate the overall discriminability among models, the Bayesian omnibus risk (BOR) was computed, reflecting the probability that all models are equally frequent in the population.

To account for residual model uncertainty and obtain robust estimates of effective connectivity, Bayesian model averaging (BMA) was applied within the PEB framework. BMA computes a weighted average of parameter estimates across the model space, with weights proportional to each model’s posterior probability (Penny et al., 2010; Stephan et al., 2010). This approach yields a principled summary of modulatory effects by integrating over competing model structures.

All analyses focused exclusively on condition-specific modulatory effects captured in the B-matrix. The B-matrix reflects how experimental conditions modulate the strength of inter-regional connections, rather than the activity of the regions. Modulatory effects are expressed in Hertz (Hz), representing changes in effective connection strength per unit time. In line with established Bayesian inference procedures, parameters with a posterior probability (PP) of being non-zero (PP ≥ 0.95) were considered statistically significant and reported in the results. As no modulation of the vmPFC → hippocampus connection reached the standard threshold in our AM condition, despite frequent reports in previous DCM studies on AM (e.g., Benoit et al., 2014; McCormick et al., 2020), we lowered the threshold to PP ≥ 0.75 to explore potential sub-threshold effects.

## 3. Results

### 3.1. Model Comparison and Selection

To identify the most plausible network configuration across the four cognitive conditions (AM, SCN, SC, OB), we performed RFX-BMS across four competing models per group (Table 1, Fig. 2).

**Table 1.** Bayesian model selection results across four experimental groups (AM, SCN, SC, OB) for four competing models. Abbreviations: AM = Autobiographical Memory; SCN = Scenarios; SC = Scene Construction; OB = Object Imagination; EPP = Expected Posterior Probability; XP = Exceedance Probability; PXP = Protected Exceedance Probability

|  | Model | EPP | XP | PXP |
| --- | --- | --- | --- | --- |
| AM | 1 | 0.59417 | 0.9209 | 0.73441 |
|  | 2 | 0.20457 | 0.0583 | 0.11159 |
|  | 3 | 0.072013 | 0.0017 | 0.07072 |
|  | 4 | 0.12924 | 0.0191 | 0.083284 |
| SCN | 1 | 0.34931 | 0.4093 | 0.27329 |
|  | 2 | 0.19513 | 0.1161 | 0.23042 |
|  | 3 | 0.10155 | 0.0239 | 0.21694 |
|  | 4 | 0.35402 | 0.4507 | 0.27934 |
| SC | 1 | 0.37316 | 0.4753 | 0.2696 |
|  | 2 | 0.27545 | 0.284 | 0.25296 |
|  | 3 | 0.13017 | 0.0512 | 0.23271 |
|  | 4 | 0.22121 | 0.1895 | 0.24474 |
| OB | 1 | 0.32514 | 0.365 | 0.25952 |
|  | 2 | 0.21802 | 0.1868 | 0.24477 |
|  | 3 | 0.31901 | 0.3821 | 0.26093 |
|  | 4 | 0.13784 | 0.0661 | 0.23478 |

**Figure 2.**
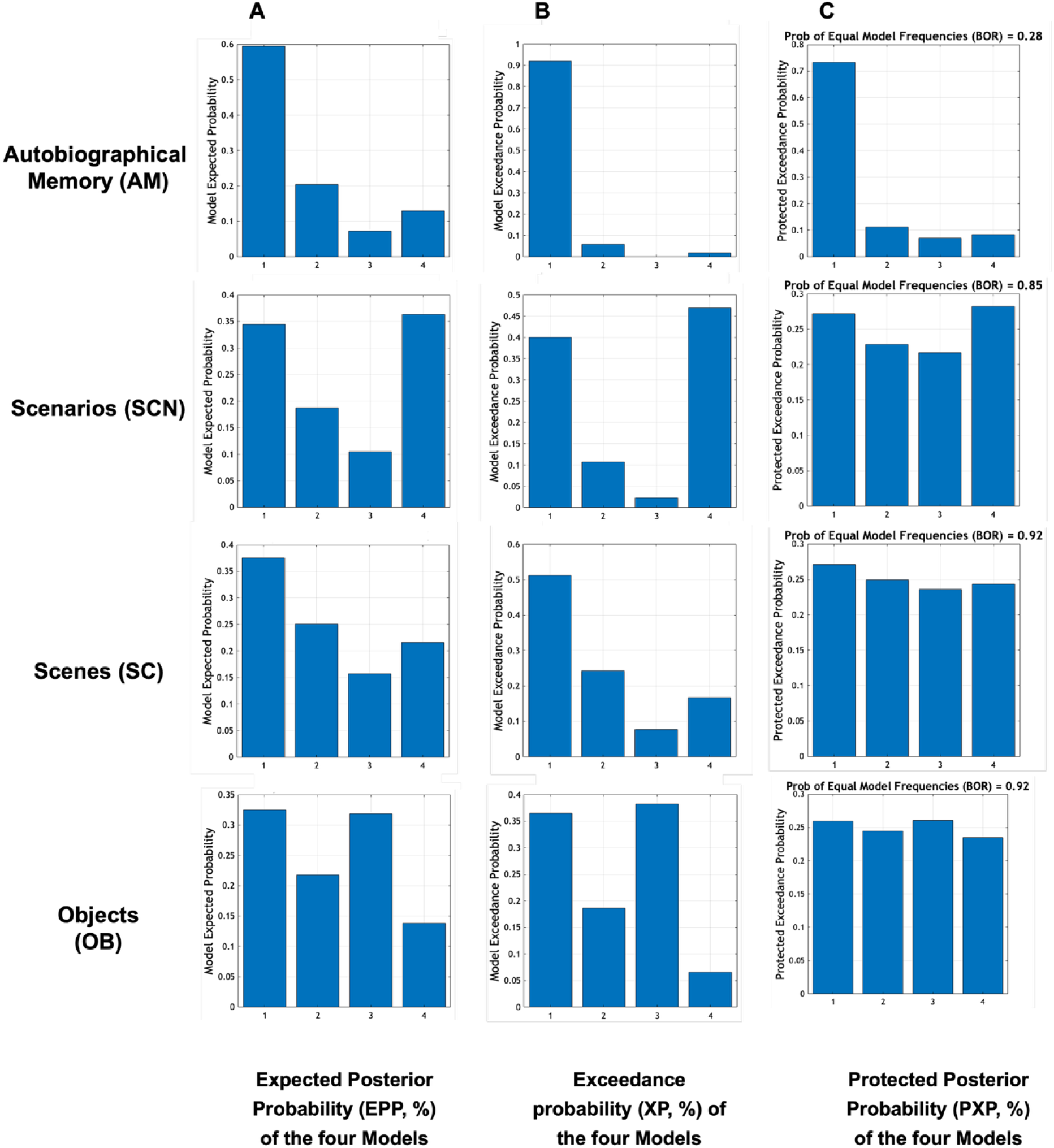
Bayesian model comparison and parameter estimation across subgroups using RFX-BMS. Model comparison is shown using the complementary metrics: expected posterior probability (A), exceedance probability (B), and protected exceedance probability with Bayesian omnibus risk (C).

In the AM condition, RFX-BMS indicated a clear preference for the first model (EPP = 0.59; XP = 0.92; PXP = 0.73; BOR = 0.28). In contrast, the SCN, SC, and OB conditions showed no clear winning model. BOR values were high (BOR ≥ 0.85), suggesting substantial inter-individual variability. Although some models showed locally increased XP values (SCN model 4: XP = 0.45; SC model 1: XP = 0.47; OB model 3: XP = 0.38), none achieved strong evidence for dominance at the group level. Our results suggest that effective connectivity during AM retrieval may follow a more consistent network pattern across individuals than during visual imagination tasks.

### 3.2. Condition-dependent B-matrix effects revealed by BMA

BMA identified condition-specific modulatory connections with PP exceeding the 0.95 threshold (Figure 3). In AM, the vmPFC showed significant negative modulation of its connections to the precuneus (B = –0.488 Hz, PP = 1.00), indicating a task-specific attenuation of prefrontal influence on parietal targets during memory retrieval. Exploratory inspection at a lowered threshold additionally revealed a modest negative vmPFC modulation on the connection to the hippocampus (B = –0.128 Hz, PP = 0.85). In SCN, the hippocampus exhibited a pronounced negative modulation of its connection to the vmPFC (B = –0.743 Hz, PP = 1.00), while its connection to the precuneus was positively modulated (B = 0.519 Hz, PP = 1.00), suggesting a divergence in hippocampal output depending on the cortical target. At the exploratory threshold, a negative modulation of the connection from the vmPFC to the hippocampus was also detected (B = –0.479 Hz, PP = 0.92). During SC, the precuneus exerted positive modulatory influences on the connection to the vmPFC (B = 0.667 Hz, PP = 1.00). No further vmPFC → hippocampus effects were observed at the lower threshold.

**Figure 3.**
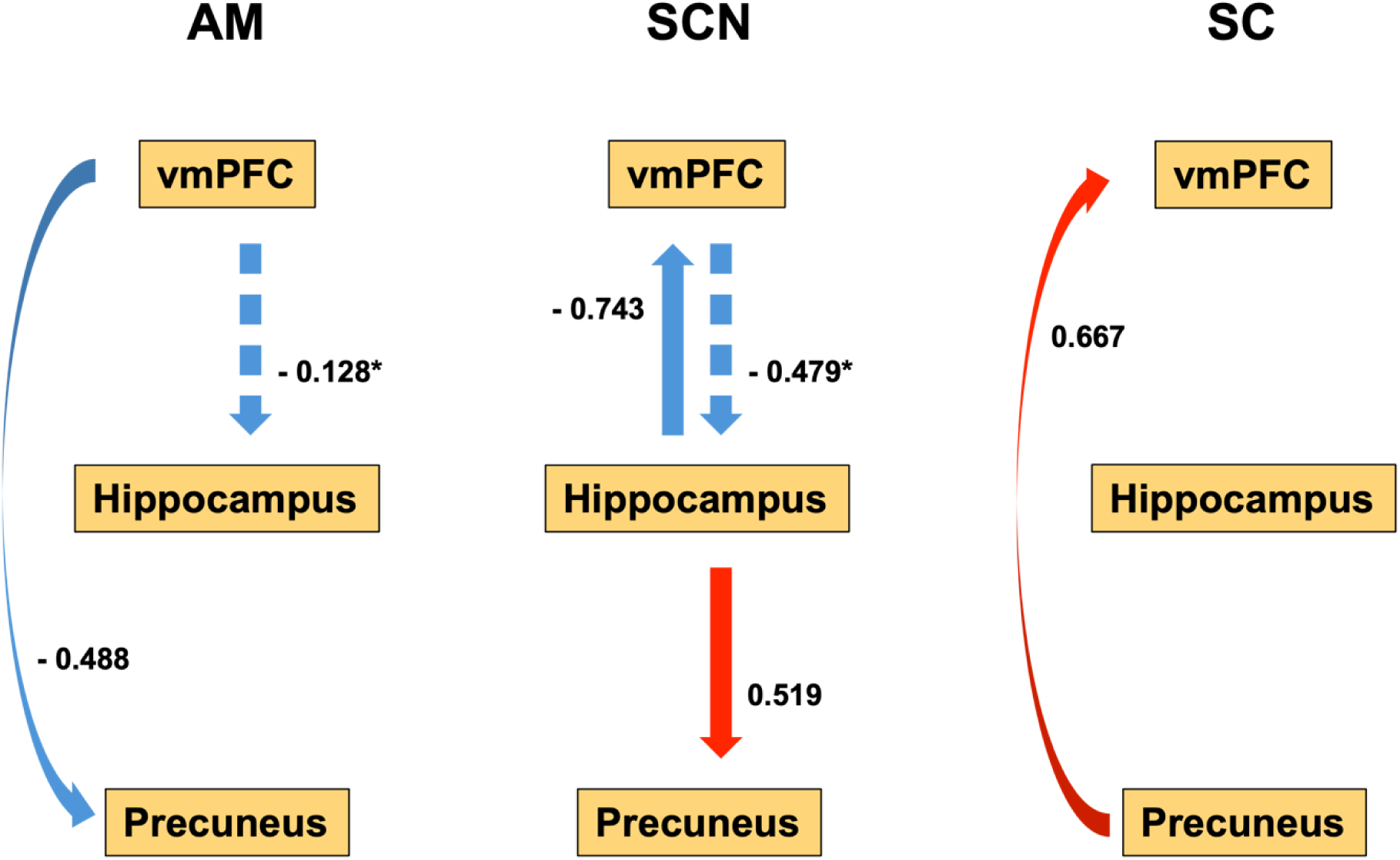
Condition-specific modulatory effects on effective connectivity (B-matrix), as revealed by BMA. Significant modulations (PP ≥ 0.95) are shown for the conditions: AM, SCN, and SC. Arrows indicate the direction and valence of modulatory effects: red represents positive modulation, blue negative modulation. Values next to each arrow indicate the estimated modulation strength in Hz, reflecting how strongly each experimental condition modulates the directed connection between two regions. Dashed arrows and asterisks next to the values denote modulatory effects with moderate evidence (PP ≥ 0.75). No modulatory connections survived the threshold in the OB imagery condition.

No significant modulatory effects were observed in the object imagery condition, and lowering the threshold likewise did not uncover a modulation of the vmPFC-hippocampus connection. Thus, while memory retrieval was associated with strong vmPFC-driven modulation, scenario and scene imagery primarily involved hippocampal and precuneus-based influences, respectively; no significant effects were observed during object imagery.

## Discussion

Using high-resolution 7T fMRI and DCM, this study investigated the directed connectivity patterns underlying AM retrieval and different forms of imaginative construction. By examining four conditions, AM, extended scenario imagination (SCN), single scene construction (SC), and object imagery (OB), we provide novel evidence for both shared and distinct directed neural dynamics between memory and imagination. In the following, we will first discuss the results concerning the overall model fit and then highlight differences between conditions in the effective connections amongst vmPFC, hippocampus, and precuneus.

### Overall Model Fit: Scene construction as the core cognitive mechanism

We found stable DCM models for three of our four conditions, namely AM, SCN, and SC. All of these conditions rely phenomenologically on the construction of naturalistic scenes (McCormick et al. 2018b; Monzel et al. 2024; Taube et al. preprint). On the contrary, OB which does not rely on scene construction (see Dalton et al. 2018) also did not result in a stable DCM model. Thus, our results highlight scene construction as a potential unifying cognitive mechanism across memory and imagination.

This pattern supports longstanding theories that the hippocampus and associated DMN regions evolved not solely for memory, but for constructive cognition more broadly—a capacity to generate spatially and temporally coherent internal representations (Hassabis et al., 2007; Maguire & Mullally, 2013). It also aligns with recent evidence that hippocampal responses scale with the naturalistic coherence of imagined scenes, rather than with image complexity per se (McCormick et al., 2021a,b).

### Autobiographical memory retrieval: vmPFC-driven effective connectivity

Among all four conditions, AM retrieval showed the most stable and coherent connectivity profile across participants. Bayesian model selection (RFX-BMS) revealed a strong preference for a model with vmPFC-driven top-down modulation, indicating robust engagement of a specific network motif during memory reconstruction. Bayesian model average (BMA) results confirmed this interpretation, revealing significant negative modulation from the vmPFC to the precuneus and, at a more exploratory threshold, to the hippocampus.

These findings reinforce the idea that vmPFC plays a central role in initiating and organizing episodic recollection, particularly for self-relevant, temporally extended events (McCormick et al., 2020; Nawa & Ando, 2019, 2020; Jacques et al., 2011; Inman et al., 2018). The negative influence on the precuneus does not necessarily reflect inhibition in a physiological sense, but rather a down-weighting or suppression of excessive visuospatial elaboration. Prior work has similarly observed vmPFC-related attenuation of posterior midline activity during memory recall (Schott et al., 2023). One possible interpretation is that vmPFC coordinates and modulates excessive spatial or self-referential elaboration, thereby maintaining coherence in complex episodic simulations. Such negative modulation might also index an internal attention mechanism, helping to regulate the balance between detailed scene visualization and overarching temporal structure. In sum, this profile underscores the vmPFC’s role as an executive hub for memory-guided simulation.

### Scenario construction: Dynamic hippocampal connectivity

In contrast to AM, the SCN condition showed no single dominant model across participants, but displayed distinct and strong modulatory effects at the connectivity level. The hippocampus negatively modulated the vmPFC and positively modulated the precuneus, with exploratory evidence of vmPFC → hippocampus modulation as well. This pattern suggests a flexible, bidirectional architecture that accommodates the complex demands of simulating temporally unfolding, novel events.

Compared to AM, scenario construction likely places greater demands on episodic generativity and real-time elaboration. The hippocampus may take on a more active role in initiating and sequencing imagined event components, while the vmPFC integrates them into a coherent temporal structure (Barry et al., 2019; Monk et al., 2020, 2021). The bidirectionality observed here contrasts with the more hierarchical pattern in AM, pointing to a network reconfiguration that emphasizes cooperation and mutual influence. This is consistent with models of future thinking that describe a distributed interplay between episodic content generation (hippocampus) and temporal control (vmPFC).

### Scene construction: Posterior midline control amid stable hippocampal engagement

The SC condition - focused on mentally generating a single, static spatial image - revealed a distinct pattern of effective connectivity: the precuneus emerged as the main driver, positively modulating the vmPFC (B = 0.667 Hz, PP = 1.00). BMS showed no clear group-level model dominance (BOR = 0.86), again indicating variability in individual strategies.

This posterior-to-anterior pattern suggests a reorientation of network dynamics away from vmPFC-led temporal orchestration (as seen in AM and SCN) toward visuospatial integration anchored in the precuneus. This finding aligns well with prior literature implicating the precuneus in self-location, perspective-taking, and spatial imagery (Svoboda et al., 2006), as well as in tasks requiring the construction of novel spatial layouts and imagined viewpoints (Hebscher et al., 2018; St. Jacques et al., 2017, 2018).

Importantly, while no significant hippocampal modulatory effects were detected in the B-matrix, this does **not** imply that the hippocampus was uninvolved in scene construction. Prior univariate analyses using the same dataset (Leelaarporn et al. 2024, Taube et al. 2025, preprint) revealed robust hippocampal activation during the SC condition, in line with a broad literature emphasizing its critical role in generating coherent spatial representations and imagined scenes (Hassabis & Maguire, 2007). The absence of dynamic hippocampal connectivity modulation in DCM may indicate that while the hippocampus is active, its influence on other nodes of the network remains tonically engaged rather than dynamically altered by the specific task demands. In this light, DCM provides a valuable complement to univariate analyses: while the latter identifies which regions are active, DCM clarifies how those regions communicate and whether the strength or direction of their influence changes across tasks. Thus, the SC findings highlight that effective connectivity can remain stable even when regional activation is strong, suggesting a division between the presence of activity and its functional regulation.

### Object construction: Minimal network reconfiguration

Finally, object construction, defined as imagining a single object without spatial or temporal context, did not reveal any significant modulatory effects, even at liberal thresholds. Model evidence was similarly inconclusive, suggesting that this task engages the vmPFC– hippocampus–precuneus circuit to a much lesser extent.

This absence of strong task-specific modulation may reflect the relatively low demands of object imagery on constructive or temporal processes. Instead, OB may primarily rely on visual working memory or ventral stream visual representations, with minimal recruitment of episodic or scene-construction systems. The lack of directed connectivity supports the interpretation that OB is cognitively and neurally simpler than the other tasks examined here.

This outcome is theoretically expected. Prior studies have associated object imagery more strongly with the perirhinal cortex than with the hippocampus or DMN regions (Lee et al. 2005; O’Neil et al., 2012). Our analysis, however, was constrained to a predefined AM/imagination network (vmPFC, hippocampus, precuneus) based on hypotheses about scene-based and episodic construction. The relative mismatch between model and task suggests that the network is not optimized for representing minimal, non-contextual object imagery.

Rather than representing a modeling failure, the instability of the object condition provides critical validation of our approach: the effective connectivity framework used here appears specifically tuned to constructive, scene-based cognition, and not broadly applicable to all forms of internal imagery. This reinforces the idea that scene construction, not simply imagery, is the common neural denominator of AM and imagination, and that the hippocampus plays a role only when spatial or temporal coherence is required.

### Toward a dynamic network model of internally generated cognition

The current results support a dynamic systems view of the default mode network, in which effective connectivity is task- and content-dependent. Rather than acting as a rigid hierarchical system, the network exhibits flexible leadership: vmPFC during temporally structured recall or simulation, precuneus during detailed spatial construction, and more distributed or unstable patterns in less structured conditions.

This view complements recent models of the DMN as a domain-general hub for internal mentation (Andrews-Hanna et al., 2014; Dijkstra et al., 2017; 2019) and fits with the concept of functional “re-use”: a single network supports multiple functions via changing patterns of directed interaction (Friston et al., 2016). Our findings contribute a connectivity-based perspective, clarifying how the brain transitions between episodic retrieval, imaginative projection, and static imagery—not just by activating shared regions, but by reconfiguring the flow of information among them. This distinction underscores the value of dynamic modeling approaches like DCM in capturing the temporal architecture of internally generated thought.

### Limitations and future directions

Several limitations should be acknowledged. First, although DCM provides rich insights into directed connectivity, we restricted our analysis to left-hemispheric ROIs for tractability. However, bilateral and other regions, such as the lateral parietal cortex, are known contributors to constructive cognition (Maguire et al., 2001; McCormick et al., 2015) and were not modeled here. Future work could incorporate more distributed or individualized network models, though this may introduce trade-offs between model complexity and interpretability.

Second, and critically, the data for AM and imagination were collected in separate studies. While acquisition protocols and preprocessing pipelines were identical, and both datasets involved 7T imaging with comparable participant groups, within-subject comparisons would offer greater inferential strength. Ideally, future studies should include all conditions in a single design, allowing for individualized DCMs and trial-by-trial modeling.

Moreover, these findings are based on healthy young adults. Clinical populations with hippocampal dysfunction—such as individuals with temporal lobe epilepsy or Alzheimer’s disease—exhibit deficits in both memory and imagination, often linked to impaired scene construction (Addis et al., 2009; McCormick et al., 2018). Future research should explore whether disrupted directional connectivity underpins these impairments, potentially revealing mechanistic targets for intervention.

## Conclusions

By comparing autobiographical memory and various forms of imagination through effective connectivity analysis, this study allows insights into how the brain dynamically constructs internal experiences. Although vmPFC, hippocampus, and precuneus consistently contribute, their directional interactions shift depending on whether one is recalling the past, simulating events, constructing scenes, or visualizing objects. Our findings highlight scene construction as the core mechanism underlying memory and imagination with temporal and spatial demands shaping network dynamics. The lack of strong hippocampal connectivity in some conditions does not imply disengagement, as univariate analyses confirm its involvement. Instead, effective connectivity reveals how information flow adapts to task demands. Together, these results support a flexible, dynamic model of constructive cognition that balances retrieval and simulation, integrates spatial and temporal elements, and balances top-down and bottom-up control across internally generated content.

## Acknowledgment and funding

We would like to thank Peter Zeidman, Xenia Kobeleva and Nadja Abdel Kafi for their valuable help and support regarding the DCM analysis. This research was supported by the Hertie Network of Excellence in Clinical Neuroscience. Work in C.M.’s lab is further financed by internal research funding of the Faculty of Medicine (BONFOR), University Hospital Bonn, and by the Deutsche Forschungsgemeinschaft (DFG, German Research Foundation, MC 244/3-1). This project is further funded by the Federal Ministry of Research, Technology and Space (BMFTR) under the funding code (FKZ): 01EO2107.

## Notes

### Competing Interest Statement

The authors have declared no competing interest.

